# Interocular Symmetry and Repeatability of Foveal Outer Nuclear Layer Thickness in Congenital Achromatopsia

**DOI:** 10.1101/367813

**Authors:** Rebecca R. Mastey, Katie M. Litts, Christopher S. Langlo, Emily J. Patterson, Margaret R. Strampe, Joseph Carroll

## Abstract

**Purpose:** To examine the intraobserver repeatability of foveal outer nuclear layer (ONL) thickness measurements and evaluate interocular symmetry for patients with achromatopsia (ACHM) and controls.

**Design:** Cross-sectional study.

**Subjects:** Sixty-four patients with *CNGA3*- or *CNGB3*-associated ACHM and 38 patients with normal vision were recruited for analysis.

**Methods:** Horizontal line scans through the fovea of each eye were acquired using optical coherence tomography. Three foveal ONL thickness measurements were made by a single observer using custom software to analyze repeatability. Interocular symmetry was assessed using the average of the three measurements for each eye.

**Main Outcome Measures:** The main parameter being measured is foveal ONL thickness.

**Results:** Mean (± SD) foveal ONL thickness for ACHM patients was 74.86 ± 17.82μm (OD) and 75.30 ± 15.68μm (OS) compared to 110.60 ± 15.67μm (OD) and 110.53 ± 13.91μm (OS) for controls. Foveal ONL thickness did not differ between eyes for ACHM (p = 0.821) or control patients (p = 0.961). Intraobserver repeatability was high for foveal ONL measurements in ACHM patients (ICC = 0.939, OD and 0.915, OS) and controls (ICC = 0.991, OD and 0.984, OS).

**Conclusions:** Foveal ONL thickness can be measured with excellent repeatability. While foveal ONL thickness is reduced in ACHM compared to controls, the high interocular symmetry indicates that contralateral ONL measurements could be used as a negative control in early-phase monocular treatment trials.

## INTRODUCTION

Congenital achromatopsia (ACHM) is an autosomal recessive inherited cone dysfunction syndrome, affecting approximately 1 in 30,000 people worldwide.^1^ It is characterized by increased light sensitivity, nystagmus, reduced visual acuity, and reduced or absent color vision. To date, mutations in six genes have been associated with ACHM: *CNGA3, CNGB3, GNAT2, PDE6C, PDE6H* and ^*ATF62–5*^— with *CNGA3* and *CNGB3* mutations accounting for nearly 70% of all ACHM cases.^6^ ACHM is generally accepted to be stable (or at worst, slowly progressing in some patients), though there are conflicting findings regarding the progressive nature of ACHM.^7–11^ The relative stability of the disease, in addition to recent success of gene therapy in the mouse, canine, and sheep^12–14^ models, has made ACHM a well-suited disease for exploring gene therapy options in humans. A number of gene therapy efforts underway are seeking to restore cone function in patients with ACHM;^15–19^ however, a prerequisite for therapeutic success is that the individual retina being treated contains residual cone photoreceptors.

A number of studies have used a variety of imaging approaches to examine remnant cone structure in patients with ACHM. Adaptive optics scanning light ophthalmoscopy (AOSLO) can be used to non-invasively image the rod and cone photoreceptors with cellular resolution.^20–22^ Confocal AOSLO imaging in patients with ACHM reveals an absence of normal waveguiding cones and an intact rod mosaic.^23–25^ In contrast, non-confocal split-detector AOSLO imaging can be used to resolve cone inner segments in a manner independent of their ability to waveguide.^26^

This technique has been used to reveal extensive, yet variable, residual cone structure in patients with ACHM.^25,26^ However, even non-confocal split-detector AOSLO-based estimates of remnant cone structure may underestimate the therapeutic potential of a given retina. This is evidenced in retinitis pigmentosa where the disease sequence starts with outer segment shortening, followed by inner segment swelling, and finally loss of the cone nuclei.^27^ Therefore, assessing the thickness of the foveal outer nuclear layer (ONL), comprised nearly exclusively of cone nuclei, may provide important auxiliary information regarding remnant cone structure in patients with ACHM.

Optical coherence tomography (OCT) has previously been used for analysis of many structures within the macular region of patients with ACHM. Variable ellipsoid zone (EZ) disruption has been observed,^10,24,25,28,29^ with the textbook presentation of a hyporeflective zone being reported between 11%-58% of cases in previous studies of ACHM.^10,24,29,30^ In addition, foveal hypoplasia (persistence of one or more inner retinal layers at the fovea) is a common feature found in patients with ACHM.^24,29,30^ With respect to the ONL, patients with ACHM show significant thinning compared to controls,^25,29,30^ though interocular symmetry has not been assessed. Defining ONL symmetry would be valuable data especially for studies where one eye is treated, and the other is used as a baseline comparison. Furthermore, the prevalence of hypoplasia can complicate delineation of the ONL boundaries in patients with ACHM, possibly affecting the repeatability of such measures. This is of great significance for studies aimed at evaluating longitudinal changes in ONL structure. Here, we used OCT to examine the interocular symmetry of the ONL at the fovea in patients with ACHM and additionally sought to establish the intraobserver repeatability of these measurements using custom software (OCT Reflectivity Analytics; ORA).^31^

## MATERIALS AND METHODS

### Subjects

This research followed the tenets of the Declaration of Helsinki and was approved by the Institutional Review Board at the Medical College of Wisconsin (PRO00030741). Images from 64 patients genetically confirmed with *CNGA3*- or *CNGB3*-associated ACHM were used for this study and 38 patients with normal vision were used as controls. The demographics of the patient populations are shown in **Table 1**. If a patient had multiple visits, the session for analysis was determined by the date that had both eyes imaged and the best image quality (assessed subjectively by a single observer, R.R.M).

**Table 1.**

| Parameter | Control | ACHM | P-value |
| --- | --- | --- | --- |
| Sex | 18M, 20F | 33M, 31F | p = 0.838 |
| Gene affected | NA | 15 *CNGA3*, 49 *CNGB3* | NA |
| Age, yrs (Mean ± SD) | 28.8 ± 10.9 | 24.8 ± 14.8 | p = 0.023* |
| **Axial Length, mm (Mean ± SD)** |  |  |  |
| OD | 24.12 ± 1.30 | 24.06 ± 2.06 | p = 0.631 |
| OS | 24.09 ± 1.31 | 24.01 ± 2.07 | p = 0.826 |

M = Male; F = Female; NA = Not applicable; \* = Statistically significant

### OCT Imaging

Prior to imaging, all patients with ACHM were dilated using either cyclomydril or a combination of tropicamide (1%) and phenylephrine hydrochloride (2.5%) for cycloplegia and pupillary dilation respectively. Control patients were imaged without dilation, as a previous study evaluated layer thickness measurements pre- and post-dilation and found no change.^32^ The Bioptigen spectral domain-OCT (Leica Microsystems, Wetzlar, Germany) was used to acquire line scans at the fovea in both eyes of each subject. Horizontal scans (1,000 A-scans per B-scan, 80–100 repeated B-scans) were obtained with a nominal scan length of 6mm or 7mm. For each eye, multiple B-scans (n = 2 to 68) were registered and averaged in ImageJ^33^ to create a single. tif image (processed line scan) with improved contrast for analysis, as previously reported.^34,35^ In two eyes from two different subjects, the line scan was not positioned at the fovea; therefore, a single B-scan was extracted from the volume scan and used for analysis. Processed line scans were cross-referenced with volume scans, when available (62 of 64 patients with ACHM), to confirm the foveal position of the line scan. For the foveal line scans, the integrity of the EZ was graded by two of three observers (K.L., C.S.L., J.C.) using previous methods.^10^ In summary, grade 1 corresponds to a continuous EZ band, grade 2 is EZ disruption, grade 3 is the absence of the EZ band, grade 4 is the appearance of a hyporeflective zone, and grade 5 is outer retinal atrophy. In cases where the two observers disagreed (19/128 eyes), the third observer graded the scan and the majority grade was used for further analysis of that scan. Three patients with grade 5 were excluded from ONL analysis as there is no outer retina present due to atrophy.

### Measuring Foveal ONL Thickness

Before quantitative ONL analysis, each processed line scan was resampled to the same scale in both the axial and lateral directions. The foveal ONL thickness was measured in the logarithmic image using a 5-pixel wide longitudinal reflectivity profile (LRP) with custom software (OCT Reflectivity Analytics; ORA).^31^ Each LRP measurement was made orthogonal to the retinal pigment epithelial layer (RPE) at the fovea, which was manually identified at the center of the deepest foveal depression. In 38 of 204 processed line scans, the image was required to be rotated as the RPE was not flat at the fovea and the LRP was unable to be rotated. This was done using Photoshop (Adobe Systems, San Jose, CA) by scaling the image so the horizontal and vertical scales matched, rotating the image relative to the RPE at the fovea, and then scaling the image back to the original dimensions. In scans with complete foveal excavation, the boundaries of the ONL were selected from the peaks of the LRP corresponding to the inner limiting membrane (ILM) and the external limiting membrane (ELM) (**Figure 1A**). In scans with foveal hypoplasia, the posterior boundary of the outer plexiform layer (OPL) was used as the anterior boundary of the ONL instead of the ILM (**Figure 1B**).^8,10^ The distance between the two peaks, representing ONL thickness, was calculated and output by the software.^31^ Three independent LRP estimates of foveal ONL thickness were obtained with the single grader (R.R.M.) masked to the prior measurements in order to assess repeatability. The mean of these three measurements was used for further analyses. Descriptive and comparative statistics were calculated using GraphPad Instat (3.1) (GraphPad Software, La Jolla, CA), with intraclass correlation coefficient (ICC) being estimated using R statistical software (R Statistical Software; Vienna, Austria). Data were tested for normality using the Shapiro-Wilk test to guide use of parametric or non-parametric statistical tests as appropriate.

**Figure 1:**
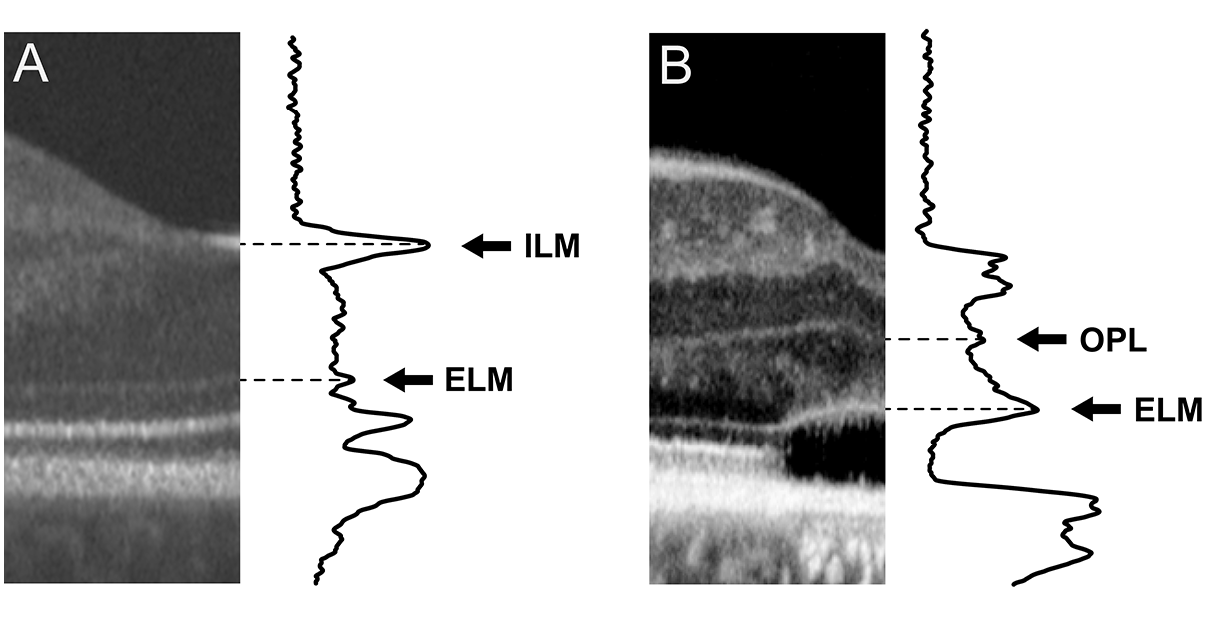
Demonstration of measuring ONL thickness in cases without hypoplasia (A) and with hypoplasia (B). When hypoplasia is not present, the ONL is defined as the distance between the ELM and the ILM. When hypoplasia is present, the ONL is then defined as the distance between the ELM and the next boundary, which is the OPL.^10,43^ Images are displayed and were measured using a logarithmic scale. ILM, inner limiting membrane. ELM, external limiting membrane. OPL, outer plexiform layer.

## RESULTS

### Subjects

As shown in **Table 1**, no statistical gender difference was observed between patients with ACHM and controls (p = 0.838, Fisher’s Exact Test) and controls were found to be significantly older than patients with ACHM (p = 0.023, Mann-Whitney Test). Axial length for patients with ACHM compared to controls was not statistically different for either eye. Furthermore, no significant interocular difference in axial length was observed for either the ACHM patients (p = 0.272, paired t-test) or controls (p = 0.824, Wilcoxon Test).

### Foveal ONL Thickness Shows High Interocular Symmetry

Consistent with previous reports,^10,25,30^ ACHM patients on average had thinner foveal ONL measurements than control patients (p < 0.0001, OD, unpaired t-test), though there was considerable overlap between the groups (**Table 2, Figure 2**). ONL thickness at the fovea did not differ between eyes for ACHM (p = 0.821, Wilcoxon test) or control patients (p = 0.961, Wilcoxon test), indicating high interocular symmetry in both populations (**Figure 2 and 3**). To assess ONL thickness in relation to EZ grade, only right eyes were used. For grade 1 scans, the mean ± SD foveal ONL thickness was 80.28 ± 16.08μm (n = 23), grade 2 scans 80.87 ± 20.60μm (n = 18), grade 3 scans 67.26 ± 12.58 μm (n = 2) and grade 4 scans 62.76 ± 10.60μm (n = 18). ONL thickness between grades 1, 2, and 4 were found to be significantly different (p = 0.0004, Kruskal-Wallis Test). There were only two patients with a grade 3, so they were not included in this comparison, and the three patients with a grade 5 were also excluded from the ONL analysis.

**Table 2.**

| Table 2. | Control |  | ACHM* |  |
| --- | --- | --- | --- | --- |
|  | OD | OS | OD | OS |
| Mean ONL Thickness (μm) | 110.60 | 110.53 | 74.86 | 75.30 |
| ± SD | ± 15.67 | ± 13.91 | ± 17.82 | ± 15.68 |
| Median ONL Thickness (μm) | 110.72 | 110.99 | 75.90 | 74.15 |
| Interquartile Range (μm) | 22.22 | 22.23 | 20.29 | 18.04 |
| ICC | 0.991 | 0.984 | 0.939 | 0.915 |
| 95% CI | 0.987-0.996 | 0.975-0.993 | 0.913-0.964 | 0.880-0.950 |
| Coefficient of Variation | 1.31% | 1.61% | 6.08% | 6.18% |

\*ICC and coefficient of variation performed on log-transformed data

**Figure 2:**
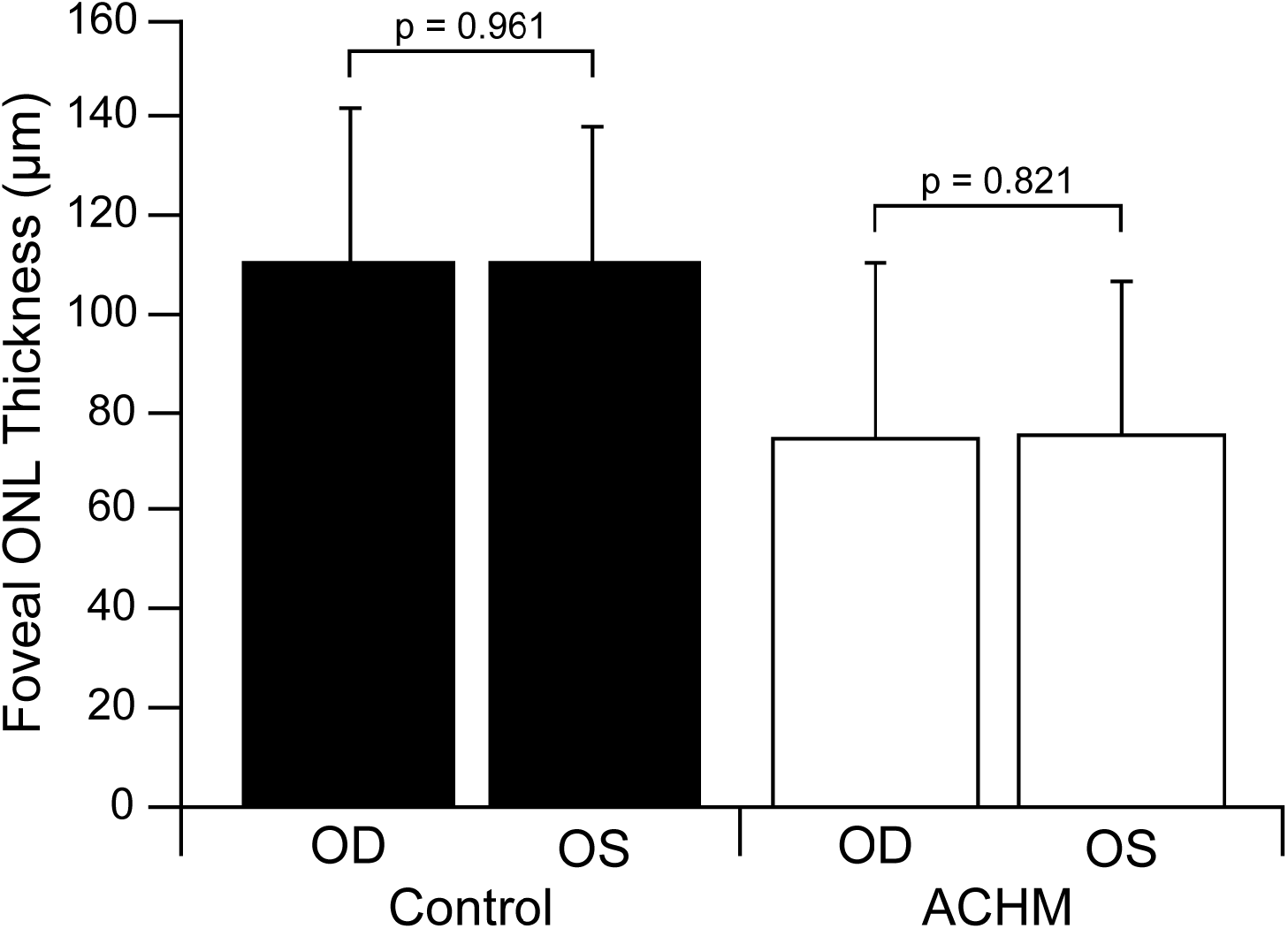
Mean foveal ONL thickness for both eyes of control and ACHM patients. On average, ACHM patients have a thinner foveal ONL than controls (p < 0.0001, OD, unpaired t-test). Error bars represent 2 standard deviations.

**Figure 3:**
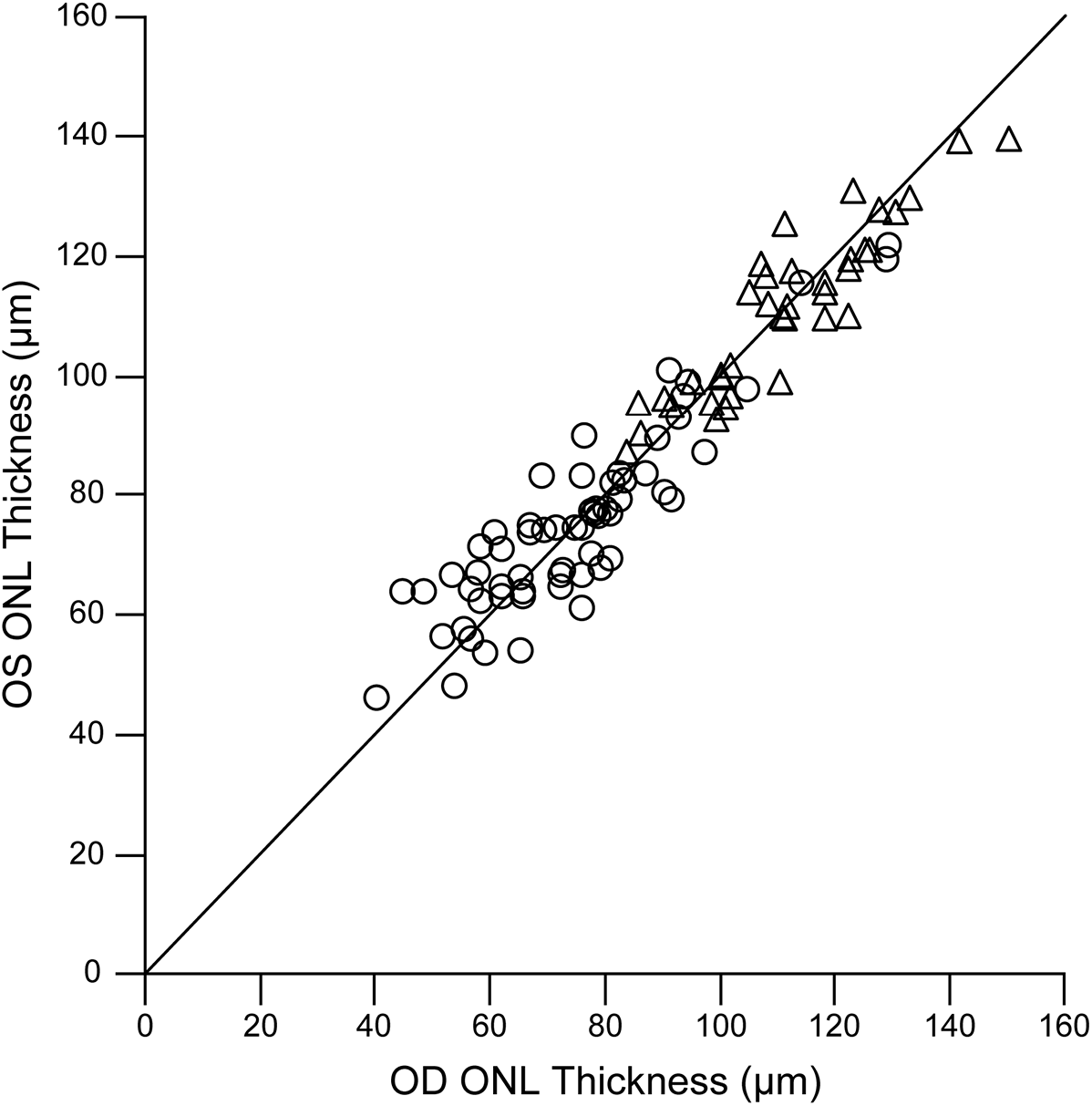
Interocular symmetry of foveal ONL thickness is shown for individuals with ACHM (circles) and controls (triangles). The solid line represents perfect interocular symmetry.

### Foveal ONL Thickness Shows Excellent Repeatability

Despite substantial variability in foveal ONL thickness within each group, excellent intraobserver repeatability was observed for our measurements. ONL data for the three measurements from controls were normally distributed, whereas data for the three measurements for ACHM patients failed normality testing. The ACHM data was log transformed, after which it passed normality—further analysis for repeatability of patients with ACHM was completed using the log transformed data. ICC values for ACHM and control patients are shown in **Table 2**. Importantly, the 95% confidence intervals for both eyes of the control patients did not overlap with the confidence intervals for the patients with ACHM. This suggests that while both groups showed excellent repeatability, the ACHM scans demonstrated slightly less repeatable than for controls. Consistent with this, measurement error, reported here as the coefficient of variation, was also higher in ACHM (**Table 2**).

## DISCUSSION

In this study we observed ONL thinning in patients with ACHM, consistent with previous studies.^10,25,29,30^ The overlap between the two groups highlights the significant variability present in ACHM. Additionally, the high repeatability of our ONL measurements suggests that the presence of foveal hypoplasia does not obviate robust assessment of ONL thickness in this challenging patient population.

Though high repeatability and ONL symmetry were observed overall, there were cases of poor repeatability in addition to examples of what appeared to be true asymmetry. Factors affecting repeatability were poor contrast and widening of boundaries as illustrated in **Figure 4**. The scan with the worst repeatability, or highest variance, was likely due to the poor contrast of the OPL in the right eye of JC_10167 (**Figure 4, *top left***). One of the three measurements was taken from the foveal reflex assuming there was no ONL, as opposed to the other measurements that were taken from the faint OPL, which was difficult to demarcate. Another possible cause for poor repeatability is the widening of the boundaries being measured, which causes the reflectivity peaks to be noisy and broad, making it hard to determine the true peak (**Figure 4, *top right***). These unclear peaks also lead to higher variation within the repeated ONL measurements. However, true asymmetry was observed in a few patients illustrated in **Figure 4** (*middle* and *bottom*). JC_10196 had the most notable asymmetry with an interocular difference of 19.54 μm, likely due to the slightly elevated ELM observed in the right eye and perhaps some residual Henle fibers at the fovea of the left eye. The subject with the second greatest interocular ONL difference was JC_10089 with a difference of 15.78 μm. The OPL in the left eye appears to be more elevated, though the signal was poor. While the overall trend supports symmetric foveal ONL thickness within patients, one needs to be aware of isolated cases of asymmetry. Likewise, image quality is a major factor influencing the repeatability of ONL thickness measurements, so the results reported here may not generalize to other studies where the overall image quality may be better or worse than that observed here.

**Figure 4:**
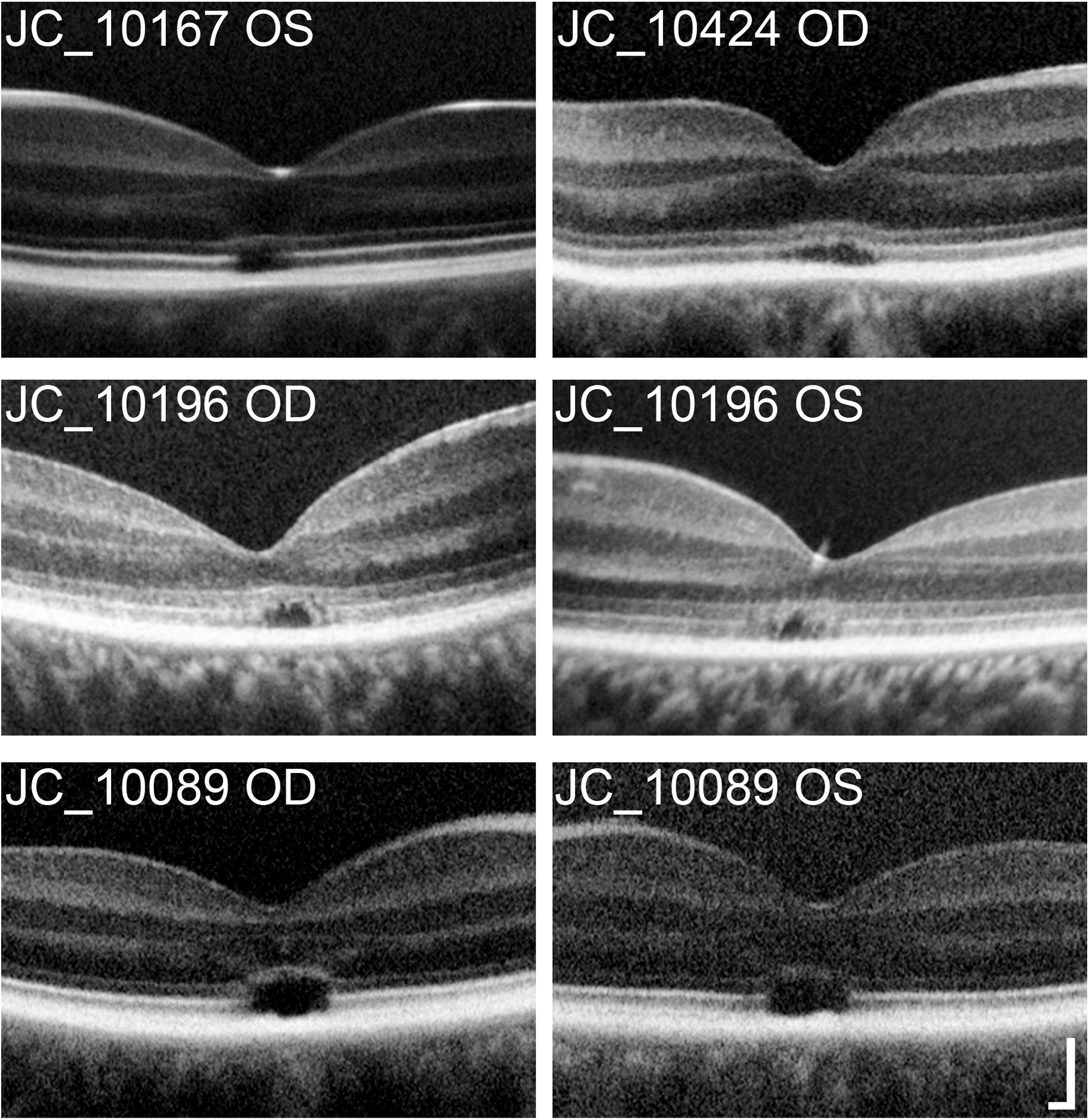
Although excellent repeatability was observed overall, there were individual eyes with poor repeatability, most commonly due to low contrast of the OPL, making demarcation difficult (*top left*) or due to imprecise, widened boundaries (*top right*). The greatest ONL asymmetry between right and left eyes was observed in JC_10196. ONL asymmetry was also observed in JC_10089; however, it is likely that low signal also played a role in this apparent asymmetry. Scale bar = 100 μm.

One limitation of our study is that the controls were significantly older than the ACHM population by an average of 4.03 years. Previous literature has conflicting reports of the effect age has on ONL measurements, with some reporting no significant changes,^36,37^ some reporting thinning,^38^ and some reporting thickening.^39,40^ The controls used in this study were significantly older than the ACHM patients; however, if decreasing ONL thickness with age is assumed, then accounting for this within our study only would have further increased our significance of ONL thickness between the two groups. On the contrary, if ONL thickness increases with age, as suggested by Chui et al., this would amount to an increase in ONL thickness of about 0.5 μm, given the 4 years age difference.^40^ As controls have a thicker ONL than ACHM by about 40 μm, the difference observed is more likely due to ACHM rather than age.

Understanding the disease progression for degenerating cone photoreceptors is also important. Milam et al. have previously observed the progression of degenerating photoreceptors in retinitis pigmentosa and reported that the shortening of the outer segment followed by the swelling of cone nuclei directly precedes complete cone photoreceptor loss.^27^

Moreover, Jacobson et al. suggested there is a staging of cone degeneration in patients with Leber Congenital Amaurosis (LCA).^41^ They proposed that within a given disease cone photoreceptors can be at varying stages of degeneration and often, regardless of intervention, some of them cannot be saved.^41^ This was observed through a LCA clinical trial that reported improved vision of treated patients, yet ONL thickness continued to decrease (which is suggestive of disease progression).^41,42^ It is currently unknown how “sick” a cone can be in ACHM and still be amenable to restoration of function via gene replacement therapy. Comprehensive assessment of the integrity of remnant foveal cones may benefit from combining ONL thickness data with measurements of cone inner segment density (assessed directly using non-confocal split-detector AOSLO) and outer segment integrity (as inferred from the relative cone reflectivity/waveguiding using confocal AOSLO).^23,25^ Such data may help to paint a more complete picture of the therapeutic potential of a given retina, which could be of use in helping frame (on a personalized basis) both patient and physician expectations regarding gene replacement therapy outcomes.

In conclusion, patients with ACHM have a thinner foveal ONL on average, though there is substantial variability between individuals. Our ONL thickness measurements derived from OCT showed both excellent intraobserver repeatability and minimal disparity in foveal ONL thickness between eyes, suggesting that the non-study eye can be used as a control in clinical trials. This is especially important as the extent of the natural progression of ACHM remains controversial.

## Acknowledgements

The authors would like to thank Brian Higgins, Alex Salmon, and Dr. Melissa Wilk for their contributions to this work.

**Financial Disclosures:** J. Carroll: MeiraGTx (Consultant), AGTC (Research Support), OptoVue (Research Support)
**Precise: (21/35 Submitted separately as ‘Highlights’ file)** We examined the intraobserver repeatability and interocular symmetry of foveal outer nuclear layer (ONL) thickness measurements in patients with congenital achromatopsia.

